# LINKER-Pred: A Deep Learning Method and Web-Server for the Prediction of Disordered Flexible Linkers in Proteins

**DOI:** 10.1101/2025.10.29.685455

**Authors:** Di Meng, Heli M. Garcia Alvarez, Juliana Glavina, César Osvaldo Leonetti, Gianluca Pollastri, Lucía Beatriz Chemes

**Affiliations:** School of Computer Science, University College Dublin, Belfield, Dublin 4, Ireland; Instituto de Investigaciones Biotecnológicas, Universidad Nacional de San Martín (UNSAM) – Consejo Nacional de Investigaciones Científicas y Técnicas (CONICET); Escuela de Bio y Nanotecnologías (EByN), Universidad Nacional de San Martín; Universidad Abierta Interamericana, Facultad de Tecnología Informática, Buenos Aires, Argentina; Universidad Abierta Interamericana, Centro de Altos Estudios en Tecnología Informática, Buenos Aires, Argentina

**Keywords:** Disordered Flexible Linker, protein language model, intrinsically disordered proteins, predictor

## Abstract

Disordered Flexible Linkers (DFLs) are unstructured regions that play critical roles in inter-domain communication and multivalent protein interactions. Despite their biological significance, the accurate identification of DFLs remains challenging due to limited experimental annotations and sparsity of dedicated prediction tools. Here we introduce LINKER-Pred, a publicly available web server featuring two convolutional neural network-based predictors trained on a novel large-scale dataset of linkers connecting folded domains (DLD dataset) and DisProt linkers. LINKER-Pred2 combines ProtTrans and MSA-Transformer embeddings within an ensemble CNN framework, achieving state-of-the-art performance on CAID2 and CAID3 benchmarks. LINKER-Pred-Lite excludes MSA-based features, improving speed while maintaining competitive predictive accuracy. LINKER-Pred predictors offer robust residue-level DFL predictions directly from sequence, providing a scalable solution for DFL annotation across proteomes. The LINKER-Pred web server and associated resources are freely available at https://linkerpred.chemeslab.org/, offering the research community an accessible tool for studying protein disorder and modularity.

## INTRODUCTION

Intrinsically disordered proteins and protein regions (collectively referred to as IDRs) carry out key cellular functions and are also closely linked to human disease [1]. IDRs are highly flexible protein regions characterized by structurally heterogeneous conformational ensembles [2]. The structural heterogeneity of IDRs and the low conservation of IDR sequences pose a significant challenge for their prediction from sequence information. One major functional class within IDRs are regions that act as disordered flexible linkers (DFLs) by connecting two folded domains and/or short linear motifs [3–6]. Tethering by DFLs mediates key functional features of proteins such as multivalency [7] and autoinhibition [8]. The annotation of DFLs within a protein sequence would improve our ability to understand IDR functions and therefore an important goal is to develop accurate predictors for DFLs.

The Critical Assessment of Protein Intrinsic Disorder (CAID) is a community-based challenge that benchmarks state-of-the-art predictors of intrinsic disorder [9–11]. Held every two years, CAID evaluates new predictors and predictors available from previous rounds. The blind test set used to benchmark predictor performance is a set of unreleased annotations from DisProt, a manually curated database of experimentally validated IDRs [12]. In addition to assessing prediction of IDRs, CAID also assesses performance on sub-tasks including functional classes such as binding regions and disordered flexible linkers. While significant progress has been made in the development of IDR predictors, prediction of binding and DFL functions faces stronger challenges. First, most top-performing predictors in the linker category of CAID2 were primarily designed for IDR prediction including SPOT-Disorder2 [13], SETH [14], and PredIDR [15], with APOD [16] being the only predictor trained specifically for DFLs that scored in the top 10. Second, predictor performance showed above-average area under the ROC curve (ROC-AUC) scores (>0.7) but average precision scores (APS) were below 0.2 for linker predictors, implying that although the models partially separate linker from non-linker residues, many high-scoring linker predictions are false positives. This highlights the struggle these models face in precisely predicting the positive class, despite moderately good overall separation between classes.

Here, we present LINKER-Pred, a family of DFL-specific prediction models trained on an expanded dataset developed by our group that includes DisProt linkers and a Domain-Linker-Domain (DLD) dataset [17]. In addition to an improved training dataset, our predictor harnesses protein language model embedding and a convolutional neural network architecture and achieves top-performance and improved APS scores when benchmarked in the CAID3 challenge. We make our predictor freely available via an easy-to-use Web Server.

## MATERIALS AND METHODS

### Training and benchmarking data

Five datasets were compiled to train and benchmark the predictors (Table S1). The DLD_IDL, DLD_IDL10, and DisProtDFL datasets were used for model training, while the CAID2Linker and CAID3Linker datasets were used in model benchmarking.

The DLD_IDL and DLD_IDL10 datasets are from the Domain-Linker-Domain (DLD) dataset [17], which comprises 2646 PDB chains, 1405 of which contain curated annotations of 1640 independent-domain linkers (IDLs) that connect two non-interacting domains [17]. IDL residues were labelled as positives (1), while all other residues were labelled as negatives (0). Accordingly, the negative class included non-linker residues from IDL-containing proteins and all residues from the 1,241 PDB chains without an annotated IDL. DLD_IDL includes all IDLs, whereas DLD_IDL10 restricts positive labels to linkers of at least ten residues, resulting in 914 annotated IDLs. Including sequences without IDLs was essential for providing sufficient negative examples for training. The DisProtDFL dataset contributes 267 DisProt sequences with 338 experimentally annotated DFLs, extending coverage to flexible linkers outside of domain boundaries and enriching the diversity of IDRs used in training [17]. In this case, all sequences contained at least one annotated DFL; all other residues were labelled as negatives. The DLD_IDL and DLD_IDL10 datasets used to train the LINKER-Pred models benchmarked in the CAID3 challenge [11] differ slightly from those used here. Here, we retrained all models using a filtered, high-quality version of the DLD dataset. The improvements include: (a) exclusion of low-resolution PDB structures (resolution >5 Å); (b) removal of short domains (<30 residues); and (c) an increased inter-domain distance threshold (Cα–Cα) of 5 Å to score interdomain contacts.

The CAID2Linker and CAID3Linker datasets, derived from the CAID2 [10] and CAID3 [11] challenges, served as external benchmarking datasets. CAID2Linker contains 40 protein sequences with 42 DFLs and was used as the *primary benchmarking set* for model optimization and comparison. CAID3Linker includes 31 protein sequences with 37 annotated DFLs and was used as an *independent benchmarking set* for external validation of generalization performance.

To prevent information leakage between training and benchmarking, we employed MMseqs2 [18] to perform sequence-similarity filtering between CAID2Linker sequences and all training-dataset sequences using the “easy-search” workflow. Training sequences with matches to CAID2Linker at either 20 % or 30% sequence identity were removed (all other parameters were left at their default values), ensuring that CAID2Linker remained fully independent during model development. This filtering step eliminated 57 training sequences in the case of the 30% sequence identity cutoff and 103 training sequences in the case of the 20% identity cutoff (Table S2). Filtering against CAID3Linker was not possible during model development, because this benchmark was released only as part of the blind CAID3 challenge and consisted of newly released DisProt linker annotations.

All datasets were labelled at the residue level, allowing the models to generate residue-wise probability scores and binary predictions for full-length protein sequences. This fine-grained approach supports high-resolution mapping of DFLs across diverse proteins.

### Overview of the prediction algorithm

We built the LINKER-Pred, LINKER-Pred Lite and LINKER-Pred2 predictors using design principles established in PUNCH2 [19]. PUNCH2 employs convolutional neural networks (CNNs) combined with modern protein language model (pLM) embeddings, particularly ProtTrans [20] and MSA-Transformer [21], to achieve high accuracy and stability in residue-level prediction of intrinsically disordered regions (IDRs). Ensemble averaging across individual models trained under 5-fold cross-validation substantially improves prediction robustness and performance [19]. Except for the refined dataset used for training (see Training and benchmarking data), the LINKER-Pred models remain identical to those benchmarked in CAID3 [11] including network architecture, pLM sequence embedding and ensemble averaging strategies.

#### Sequence representation

Two pre-trained protein language models were employed for sequence representation: ProtTrans [20] captures intrinsic sequence-level information directly from a large sequence data set from UniProt, while MSA-Transformer [21] encodes evolutionary and co-variation patterns derived from multiple sequence alignments (MSAs). Although MSA-Transformer embeddings are more computationally intensive, they provide richer evolutionary context, complementing ProtTrans’s intrinsic sequence-based representations. Each amino acid residue is represented as a high-dimensional vector (1024-dimensional for ProtTrans and 768-dimensional for MSA-Transformer), producing an embedding matrix of size *N* × *D* for a protein of length *N*. For one-dimensional convolution, this matrix is arranged with the *D* embedding features as input channels and the *N* residues along the sequence-length dimension. In addition, one-hot encoding was used as a simpler baseline representation, in which each residue is encoded as a binary vector indicating amino acid identity, with a single non-zero value corresponding to the observed residue type.

#### Network architecture

We adopted a neural network termed M2M (Many-to-Many), composed exclusively of one-dimensional convolutional layers, as the core learning architecture (Figure 1A). Compared with the deeper architectures explored in PUNCH2 [19], M2M is smaller and shallower, suitable for the reduced dataset size used here. M2M consists of three convolutional layers: the first two (conv_1_, conv_2_) apply ReLU activation functions to transform the embedded sequences and extract local contextual features, and the final layer (conv_last_) projects these representations into a one-dimensional vector matching the sequence length. Conv_1_ uses a kernel size of *K_1_* = 1 and maps the *D* input feature channels to *C_1_* = 50 output channels. Thus, it performs a learned position-wise projection of each residue embedding from 1,024 or 768 dimensions to 50 features. Conv_2_ uses *K_2_* = 5 and *C_2_* = 5, integrating information across neighbouring residues, whereas conv_last_ uses *K_last_* = 1 and *C_last_* = 1 to generate one output score per residue. This design enables position-wise residue classification with low computational cost. The total number of parameters depends on the embedding dimensionality as this determines the input channels of the first convolutional layer. Since convolutional filters are shared across sequence positions, the number of trainable parameters does not depend on protein length.

**Figure 1.**
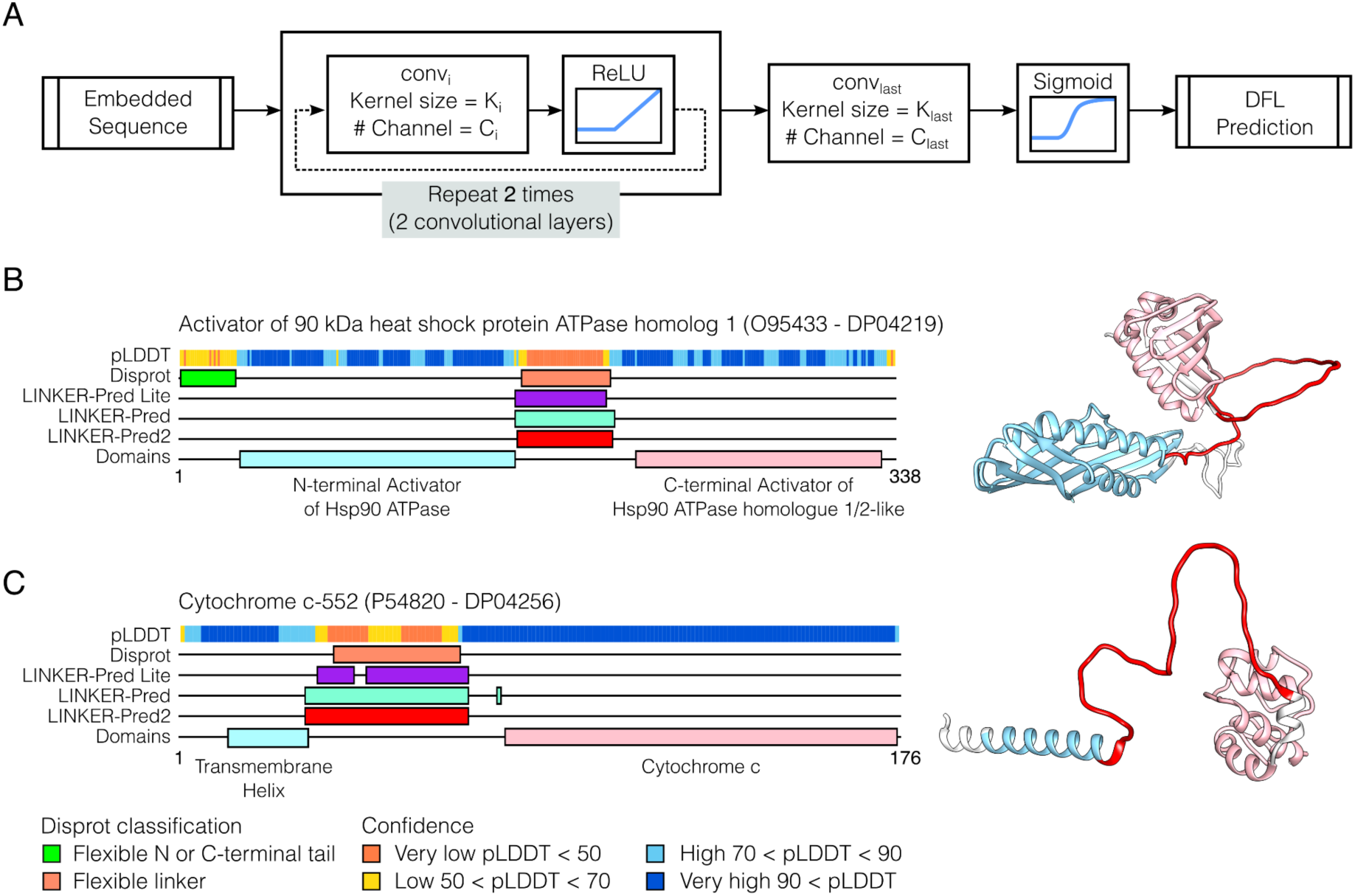
Architecture of LINKER-Pred models and example predictions. (A) The (M2M) neural network architecture consists of two intermediate (conv_i_) and one output (conv_last_) convolutional layers. B-C) Schematic representation of the Activator of 90 kDA heat shock protein ATPase homolog 1 (B) and the Chromobox protein homolog 5 (C). From top to bottom: AlphaFold2 [22] pLDDT score, DisProt annotations for Disorder Function, LINKER-Pred predictions and Pfam domain or structural elements annotations. Right: AlphaFold2 models with LINKER-Pred2 prediction marked in red.

#### Ensemble strategy

Rather than concatenating embeddings from different models into a single high-dimensional input, we applied an ensemble learning strategy, training separate CNNs with identical M2M architecture for each embedding type. Multiple models were trained using 5-fold cross-validation to reduce variance and to enhance generalization. The resulting ensembles were aggregated by averaging their residue-level probability outputs. We developed three predictors differing in ensemble composition: LINKER-Pred-Lite, LINKER-Pred, and LINKER-Pred2 (Table S3). LINKER-Pred-Lite (5 models) comprises five ProtTrans-based models trained with five-fold cross-validation, LINKER-Pred (10 models) adds five additional MSA-Transformer-based models, and LINKER-Pred2 (15 models) extends the ensemble with further MSA-Transformer models trained on additional datasets such as DLD_IDL10 which filters out short (<10 residues) linkers.

### Training data partitioning and individual CNN model training

After removing training sequences with similarity to the CAID2Linker benchmark dataset, the remaining sequences were partitioned into five folds for model training. To construct these partitions, the retained training sequences were first clustered using MMseqs2 [18] “easy-cluster”, applying the same sequence identity cutoffs used during CAID2Linker filtering, together with an 80% coverage threshold (-c 0.8) in coverage mode 1 (--cov mode 1) (all other parameters were left at their default values). Thus, datasets filtered against CAID2Linker at 30% sequence identity were subsequently clustered at 30% identity, whereas datasets filtered at 20% identity were clustered at 20% identity. The resulting independent sequence clusters were then assigned to five non-overlapping groups, ensuring that sequences within the same similarity cluster were kept in the same fold. Each fold was used once as the validation set while the remaining four folds were used for training, yielding five individual CNN models for each dataset–embedding configuration (Table S3).

As an additional control, we also generated a random five-fold partition of the training sequences retained after 20% sequence identity filtering against CAID2Linker, providing a comparison with the sequence identity-based partitioning strategy. Table S4 summarizes the composition of the corresponding final combined training sets after CAID2Linker similarity filtering, including the number of retained sequences, positive and negative sequences, and number of annotated DFL regions.

Individual CNN models were trained for a fixed number of epochs without early stopping. One-hot- and ProtTrans-based models were trained for 120 epochs for all dataset combinations. MSA-Transformer-based models trained on DLD_IDL + DisProtDFL were trained for 150 epochs, whereas MSA-Transformer-based models trained on DLD_IDL10 + DisProtDFL were trained for 120 epochs.

### Model performance evaluation

For each individual CNN model described above, training behaviour was first assessed by inspecting training and validation loss curves, together with validation ROC-AUC (Area Under the Receiver-Operating Characteristic Curve) across epochs. Final ensemble predictive performance was then evaluated using cross-validation and independent benchmark datasets.

For cross-validation, predictions for each held-out fold were generated only using models trained on the remaining four folds. For ensemble predictors, residue-level probability outputs were averaged across the corresponding models that had not been trained on the held-out fold. Global out-of-fold metrics were then computed by pooling these held-out residue-level predictions across the five folds, whereas per-fold metrics were computed separately to assess variability across folds. Performance was quantified using the ROC-AUC, the APS (Average Precision Score), and the maximum F1 score across decision thresholds (F1 max).

Independent benchmarking was performed on CAID2Linker and CAID3Linker. For both benchmark datasets, final ensemble predictions were obtained by averaging the residue-level probability outputs of all individual models included in each predictor, and then computing the corresponding ROC-AUC, APS, and F1 max metrics.

### Classification of AlphaFold-defined regions

AlphaFold models [22] of proteins in the CAID3Linker dataset were classified into folded and disordered regions using a procedure adapted from Tesei et al. (2024) [23]. Per-residue pLDDT scores were rescaled from 0–100 to 0–1. Residues with pLDDT > 0.8 were initially classified as folded, residues with pLDDT < 0.7 as disordered, and residues with 0.7 ≤ pLDDT ≤ 0.8 as gaps. Contiguous folded or disordered segments shorter than 10 residues were reassigned as gaps. Gap segments were subsequently classified as disordered when flanked by disordered segments on both sides or when located at a sequence terminus adjacent to a disordered segment; all remaining gaps were classified as folded. The resulting disordered regions were categorized according to their sequence position. Terminal regions were labelled as “N-terminal” or “C-terminal. Internal disordered regions were labelled as “Inter-domain linker” when they were flanked on both sides by folded segments of at least 30 residues. All other internal disordered regions, for which at least one flanking folded segment was shorter than 30 residues, were classified as “Other.”

## RESULTS AND DISCUSSION

### Cross-validation performance

We first evaluated the LINKER-Pred family predictors using five-fold cross-validation under progressively more stringent sequence-partitioning schemes (Table 1). Random partitioning was included as an internal reference setting, whereas MMseqs2 cluster-based partitioning at 30% and 20% sequence identity cutoffs was used to reduce similarity between training and validation folds and obtain more conservative estimates of generalization performance. As expected, methods trained with random partitioning produced the highest performance estimates. In contrast, APS and F1 max decreased modestly for methods trained under cluster-based partitioning, with the lowest values observed at the 20% sequence-identity cutoff, while ROC-AUC remained high for all three ensemble predictors. Importantly, method performance did not markedly decrease under cluster-based partitioning. This is consistent with the high sequence diversity of the training set. Clustering the original 2,913 training set sequences yielded 1,759 clusters at 30% sequence identity and 1,571 clusters at 20% sequence identity; singleton clusters accounted for 68.7% and 63.5% of all clusters, respectively. Thus, the predictors retained robust performance even when cross-validation folds were defined by sequence-identity clustering.

**Table 1.**
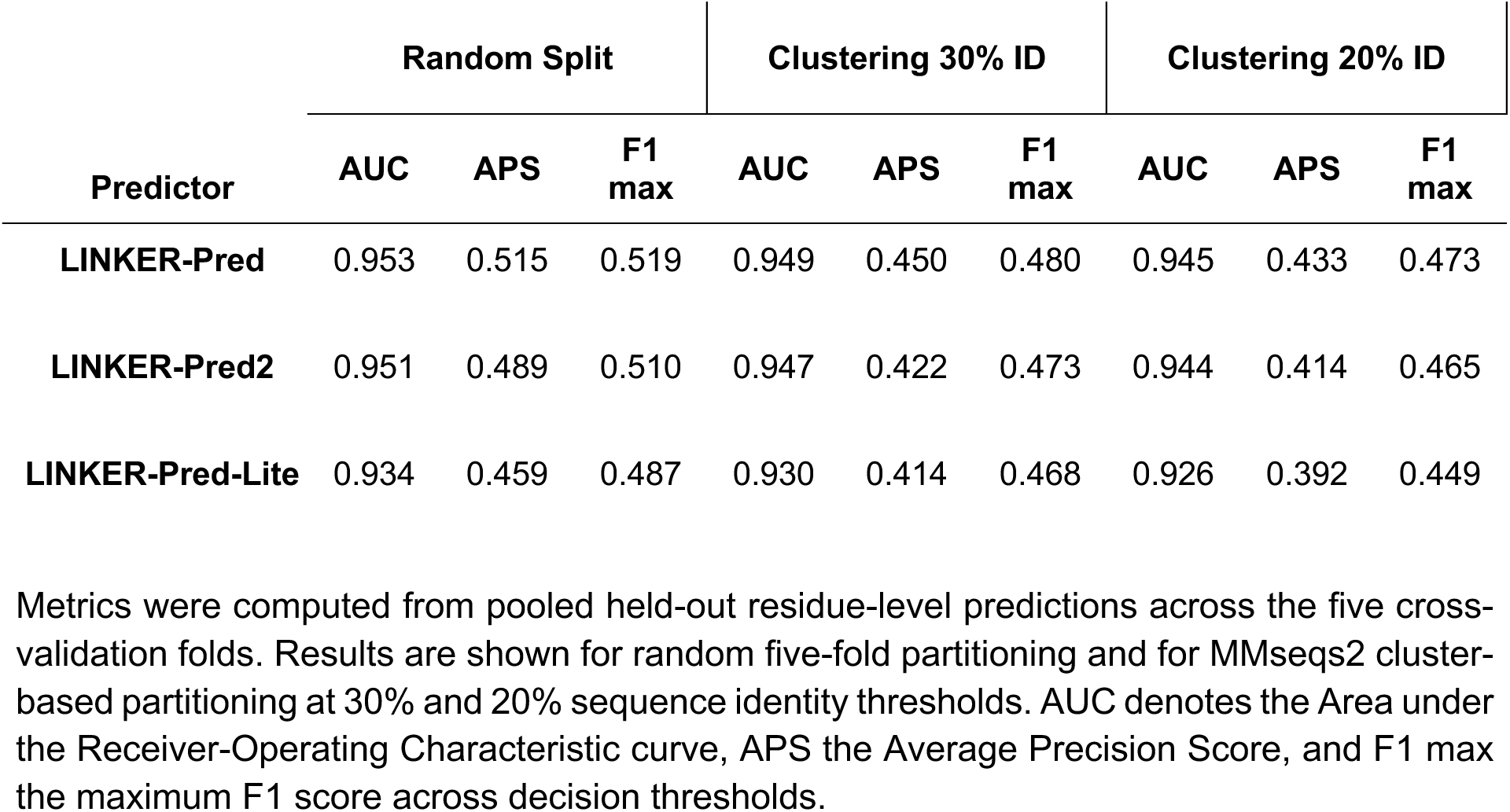
Cross-validation performance of LINKER-Pred family predictors under different training dataset partitioning schemes.

| Predictor | Random Split |  |  | Clustering 30% ID |  |  | Clustering 20% ID |  |  |
| --- | --- | --- | --- | --- | --- | --- | --- | --- | --- |
|  | AUC | APS | F1 max | AUC | APS | F1 max | AUC | APS | F1 max |
| LINKER-Pred | 0.953 | 0.515 | 0.519 | 0.949 | 0.450 | 0.480 | 0.945 | 0.433 | 0.473 |
| LINKER-Pred2 | 0.951 | 0.489 | 0.510 | 0.947 | 0.422 | 0.473 | 0.944 | 0.414 | 0.465 |
| LINKER-Pred-Lite | 0.934 | 0.459 | 0.487 | 0.930 | 0.414 | 0.468 | 0.926 | 0.392 | 0.449 |

Metrics were computed from pooled held-out residue-level predictions across the five cross- validation folds. Results are shown for random five-fold partitioning and for MMseqs2 cluster- based partitioning at 30% and 20% sequence identity thresholds. AUC denotes the Area under the Receiver-Operating Characteristic curve, APS the Average Precision Score, and F1 max the maximum F1 score across decision thresholds.

We therefore used the 20% identity cluster-based split, the most stringent validation setting, to further inspect the training behaviour of the underlying individual CNN models. Across folds and training epochs, training and validation loss curves showed no systematic divergence, indicating that model optimization did not result in severe overfitting. Validation ROC-AUC also increased during training before reaching a plateau, consistent with improved validation performance without a subsequent decline (Figure S1). As an additional control, we trained equivalent CNN models using one-hot encoding instead of protein language model embeddings. These models reached validation ROC-AUC values of approximately 0.65, substantially below the embedding-based models, indicating that the learned sequence representations play a central role in LINKER-Pred performance. Per-fold ensemble metrics were also consistent across validation folds for all partitioning schemes (Figure S2), supporting the robustness of the global out-of-fold cross-validation metrics reported in Table 1. Accordingly, all subsequent analyses refer to predictors trained using the 20% sequence-identity cluster-based partitioning scheme.

### Independent benchmark performance

LINKER-Pred predicted DisProt-annotated DFLs including linkers connecting folded domains (Figure 1B) and other linkers such as those tethering domains and transmembrane helices (Figure 1C). The original LINKER-Pred models achieved top-performance amongst DFL-specific predictors in CAID3 [11]. Next, we benchmark the predictors retrained on the refined DLD dataset against top-performing DFL prediction methods from CAID2 and CAID3 [10,11] (Table 2).

**Table 2.**
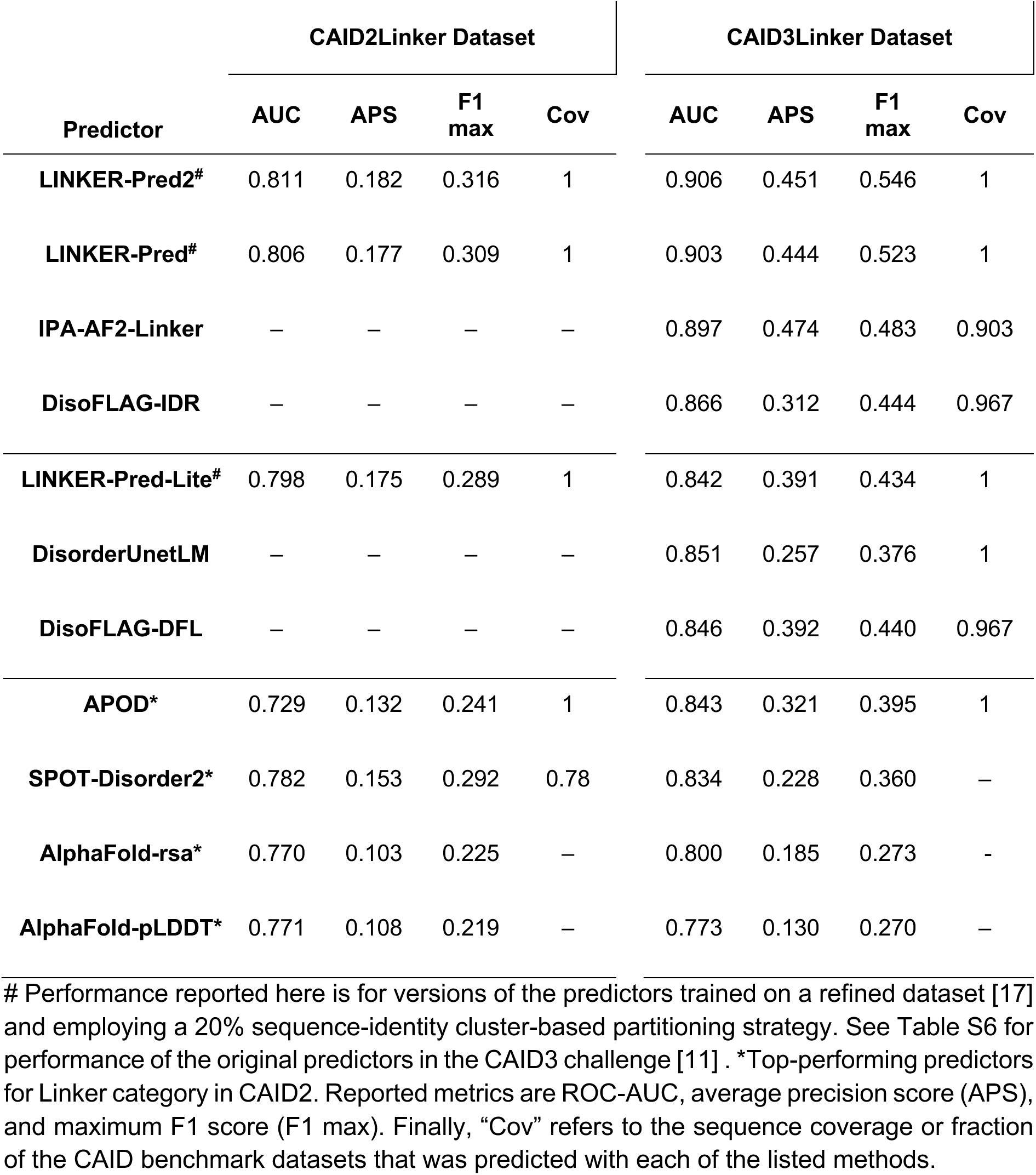
Benchmarking of LINKER-Pred family predictors on the CAID2 and CAID 3 datasets.

| Predictor | CAID2Linker Dataset |  |  |  | CAID3Linker Dataset |  |  |  |
| --- | --- | --- | --- | --- | --- | --- | --- | --- |
|  | AUC | APS | F1 max | Cov | AUC | APS | F1 max | Cov |
| **LINKER-Pred2<sup>#</sup>** | 0.811 | 0.182 | 0.316 | 1 | 0.906 | 0.451 | 0.546 | 1 |
| **LINKER-Pred<sup>#</sup>** | 0.806 | 0.177 | 0.309 | 1 | 0.903 | 0.444 | 0.523 | 1 |
| **IPA-AF2-Linker** | – | – | – | – | 0.897 | 0.474 | 0.483 | 0.903 |
| **DisoFLAG-IDR** | – | – | – | – | 0.866 | 0.312 | 0.444 | 0.967 |
| **LINKER-Pred-Lite<sup>#</sup>** | 0.798 | 0.175 | 0.289 | 1 | 0.842 | 0.391 | 0.434 | 1 |
| **DisorderUnetLM** | – | – | – | – | 0.851 | 0.257 | 0.376 | 1 |
| **DisoFLAG-DFL** | – | – | – | – | 0.846 | 0.392 | 0.440 | 0.967 |
| **APOD<sup>\*</sup>** | 0.729 | 0.132 | 0.241 | 1 | 0.843 | 0.321 | 0.395 | 1 |
| **SPOT-Disorder2<sup>\*</sup>** | 0.782 | 0.153 | 0.292 | 0.78 | 0.834 | 0.228 | 0.360 | – |
| **AlphaFold-rsa<sup>\*</sup>** | 0.770 | 0.103 | 0.225 | – | 0.800 | 0.185 | 0.273 | – |
| **AlphaFold-pLDDT<sup>\*</sup>** | 0.771 | 0.108 | 0.219 | – | 0.773 | 0.130 | 0.270 | – |

<sup>#</sup> Performance reported here is for versions of the predictors trained on a refined dataset [17] and employing a 20% sequence-identity cluster-based partitioning strategy. See Table S6 for performance of the original predictors in the CAID3 challenge [11]. <sup>\*</sup>Top-performing predictors for Linker category in CAID2. Reported metrics are ROC-AUC, average precision score (APS), and maximum F1 score (F1 max). Finally, “Cov” refers to the sequence coverage or fraction of the CAID benchmark datasets that was predicted with each of the listed methods.

### Performance on CAID2 (primary benchmarking)

On the CAID2Linker benchmark, all LINKER-Pred models outperformed the top predictors across AUC and APS metrics (Table 2). LINKER-Pred2 achieved the highest AUC (0.811), followed by LINKER-Pred (0.806) and LINKER-Pred-Lite (0.798). APS values for LINKER-Pred models (0.175-0.182) were higher than the top CAID2 predictors, including SPOT-Disorder2 (AUC = 0.782, APS = 0.153) and average AlphaFold-derived scores (AUC ≈ 0.771, APS ≈ 0.106). Most predictors in the CAID2Linker benchmark were trained for general IDR prediction. The only DFL-specific predictor, APOD, performed substantially lower (AUC = 0.729, APS = 0.132). All LINKER-Pred models achieved 100% coverage, successfully generating predictions for every sequence in the dataset, an advantage over several CAID2 predictors that failed on some test proteins.

### Performance on CAID3 (independent benchmarking)

The CAID3Linker dataset served as a fully independent test set, offering the most stringent assessment of model generalization. The original LINKER-Pred methods scored as top-performers in the CAID3 Linker challenge, with LINKER-Pred2 scoring second best below IPA-AF2-Linker (Table S6) [11,24]. The refined LINKER-Pred2 and LINKER-Pred models achieved the strongest overall performance among the evaluated methods (Table 2). LINKER-Pred2 obtained the highest ROC-AUC (0.906) and F1 max (0.546), followed closely by LINKER-Pred (ROC-AUC, 0.903; F1 max, 0.523). Both methods ranked second in average precision score (APS), with values of 0.451 and 0.444, respectively. LINKER-Pred-Lite also performed competitively (AUC = 0.842, APS = 0.391 and F1 max = 0.434), providing a favourable trade-off between speed and predictive performance.

Because F1 max is obtained by evaluating F1 across classification thresholds, the corresponding optimal threshold is dataset dependent. On CAID3Linker, the maximum F1 scores for LINKER-Pred2, LINKER-Pred, and LINKER-Pred-Lite were obtained at thresholds of 0.130, 0.110, and 0.150, respectively; the corresponding CAID2Linker values are reported in Table S5. These thresholds were determined following the CAID3 evaluation methodology [11], and should be interpreted as dataset-specific operating points rather than as intrinsic properties of the predictors. In contrast, APS summarizes performance across the full precision–recall curve and therefore provides a threshold-independent measure of a method’s ability to prioritize linker residues.

The refined LINKER-Pred and LINKER-Pred2 methods (Table 2) outperformed the original models (Table S6) supporting the value of the implemented data filtering strategy, and despite having been developed using a more stringent sequence-identity-based training data partitioning scheme. Overall, the performance of the LINKER-Pred models is comparable to or better than the leading CAID3 predictors [11], including IPA-AF2-Linker, DisoFLAG-IDR, and DisoFLAG-DFL (Table 2) and LINKER-Pred methods again achieved 100% coverage, underscoring their reliability and scalability.

Overall, the LINKER-Pred models demonstrated competitive AUC and APS scores compared to predictors across both benchmarks. The performance on the CAID3 dataset indicates strong generalization across datasets and linker types. These results establish LINKER-Pred methods as state-of-the-art predictors for residue-level DFL identification.

### Assessment of region-specific predictions on CAID3

To further evaluate the regional specificity of the LINKER-Pred methods, we examined predictions across DisProt-annotated DFLs and AlphaFold pLDDT-defined low-confidence regions in CAID3Linker proteins (full-length proteins containing linkers from the CAID3Linker dataset), distinguishing “Inter-domain linker” (IDL) regions from “N- or C- Terminal” regions and from “Other” internal low-confidence regions that did not satisfy the “Inter-domain linker” definition (see Materials and Methods for details on region definition). This analysis assessed whether the models preferentially identified DisProt-annotated DFLs or IDL regions rather than broadly predicting disordered or low-confidence segments.

Across CAID3Linker proteins, all three methods preferentially predicted DisProt-annotated DFLs and pLDDT-defined IDLs, although the fraction of residues predicted as linker varied considerably among individual regions (Figure S3A). Predictions within DisProt-annotated and pLDDT-derived N- and C-terminal regions were generally negligible, whereas regions classified as “Other” showed intermediate and more variable prediction levels, particularly for LINKER-Pred and LINKER-Pred2.

To determine to what extent pLDDT-defined regions coincided with experimental linker annotations, we stratified these regions according to whether they overlapped a DisProt-annotated DFL by at least one residue (Figure S3B). For all three models, IDLs with DisProt overlap showed higher predicted linker fractions than those without overlap. A similar enrichment was observed for regions classified as “Other”, particularly for LINKER-Pred and LINKER-Pred2, whereas N-terminal regions remained weakly predicted.

Since this binary classification does not reflect the extent of overlap, we next quantified, within overlapping IDLs, the fraction of each region covered by the DisProt annotation and by each LINKER-Pred prediction (Figure S3C). LINKER-Pred and LINKER-Pred2 frequently covered a larger fraction of the region than the corresponding DisProt annotation, suggesting that the models may identify candidate linker residues beyond the experimentally annotated segment. Although these results are promising, this possibility should be assessed in a larger dataset.

Together, these results support the regional specificity of the LINKER-Pred methods and suggest that they may extend experimentally annotated linker boundaries within structurally plausible IDL regions. This hypothesis should nevertheless be evaluated in a larger dataset combining experimentally curated linker annotations with structural annotations derived from available AlphaFold models across the human proteome.

## SERVER DESCRIPTION

### Overview of the website framework

The LINKER-Pred web server is an integrated web platform for predicting and visualizing disordered flexible linkers (DFLs) in proteins. It hosts two complementary predictors, LINKER-Pred-Lite and LINKER-Pred2, both trained using the 20% sequence-identity cluster-based partitioning scheme. LINKER-Pred-Lite is designed for rapid, multi-sequence throughput, while LINKER-Pred2 is intended for single-sequence, precision-focused analyses which requires the independent generation of an input MSA file.

Although LINKER-Pred showed slightly higher cross-validation metrics, LINKER-Pred2 was selected for the web server because it incorporates models trained on the DLD_IDL10 dataset, which applies a more stringent biological definition of linkers by restricting positive labels to IDLs of at least ten residues, as also currently implemented in the DisProt database. This criterion reduces the contribution of very short segments, while imposing only a minimal increase in computational cost. Consistent with this rationale, LINKER-Pred2 achieved the highest AUC and APS values on both the CAID2Linker and CAID3Linker independent benchmark datasets.

The two selected methods provide a practical balance between top predictive performance and computational efficiency for real-world use. Further details on the backend architecture, implementation, computational requirements, and runtime performance of the web server are provided in the Supplementary Note.

### Web Pages

The web interface of the LINKER-Pred web server is designed to be clean, intuitive, and functionally focused (Figure 2). The Download button located at the top of the Home page (Figure 2A, label 1) provides access to the training and validation datasets and to source code through ready-to-use Docker container images or a GitHub repository, enabling users to integrate LINKER-Pred tools into larger workflows. The Help button (Figure 2A, label 2) provides a brief description of the predictors, and a description of the results file.

**Figure 2.**
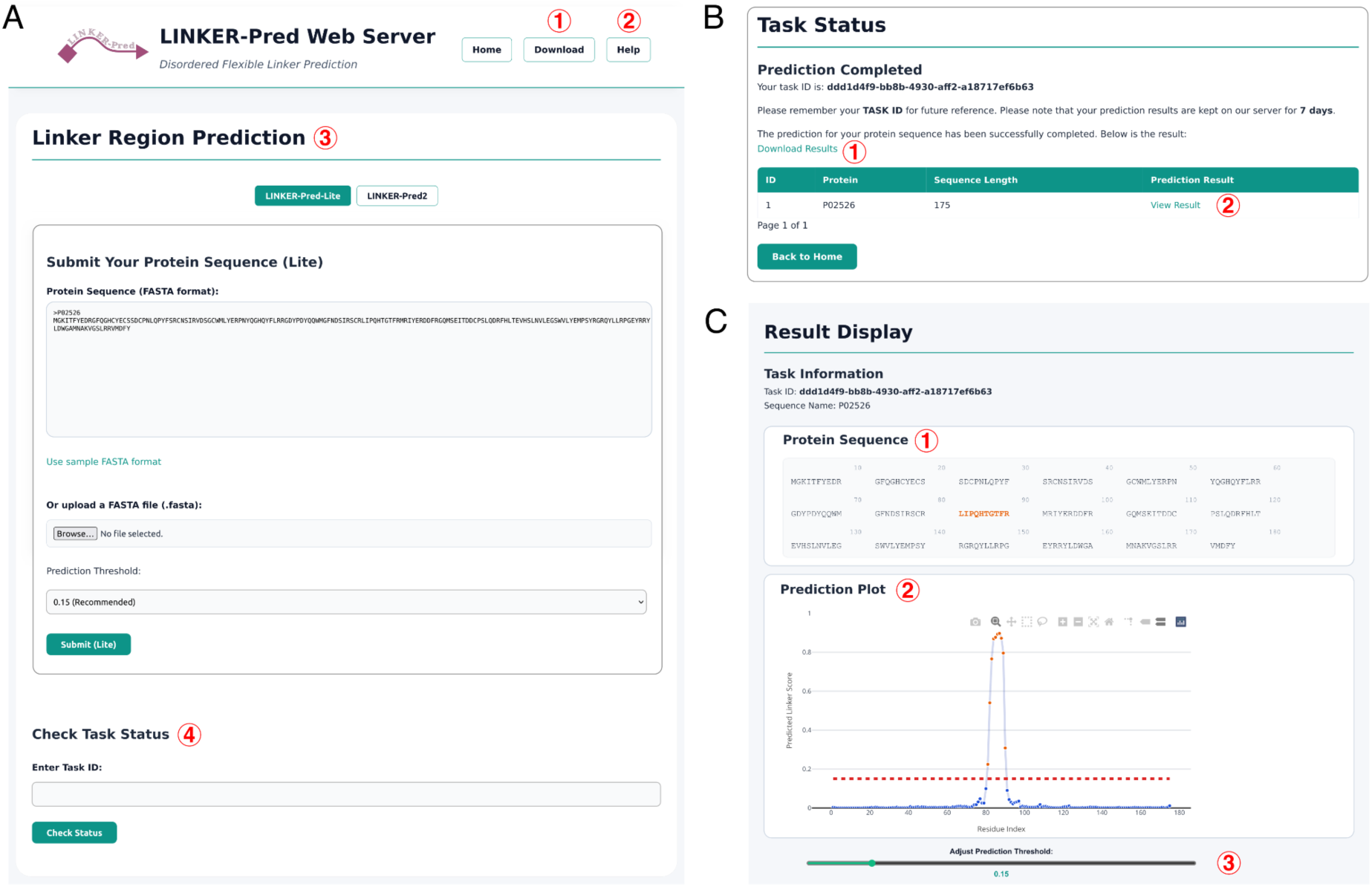
LINKER-Pred Web Server. (A) LINKER-Pred web server homepage. (B) Task Status page for a completed task. (C) Results Display page. The region with a score above the selected threshold is highlighted in red in the sequence.

The Home page contains the “Linker Region Prediction” panel (Figure 2A, label 3), which displays two tabs that allow users to switch easily between LINKER-Pred-Lite and LINKER-Pred2. LINKER-Pred-Lite requires only the sequence input in FASTA format, while LINKER-Pred2 requires the sequence plus an MSA file in .a3m format which must be generated independently using the HHblits web server [25]. Each submission generates a unique Task ID and redirects the user to the “Task Status” page (Figure 2B), which displays real-time updates on job progress. Once complete, the page provides links to download (Figure 2B, label 1) and view (Figure 2B, label 2) the results. Clicking the “View Results” link opens the “Result Display” page (Figure 2C) which displays the protein sequence and an interactive scatter plot that displays per-residue scores (range 0-1). Users can dynamically adjust the prediction threshold to explore the precision-recall trade-off and refine linker-boundary resolution (0.15 is provided only as a practical suggested operating point). Completed jobs are stored for seven days and can be retrieved from the “Check Task Status” section using the Task ID (Figure 2A, label 4).

### User guide and example

As an example, UniProt P02526 can be submitted through the LINKER-Pred-Lite tab by pasting the FASTA header and sequence, selecting the desired threshold (default: 0.15), and submitting the job (Figure 2A, label 3). After submission, the system validates the input format, launches the job, and redirects the user to the “Task Status” page (Figure 2B). When the job is complete, users can download the results as a ZIP archive or open the interactive result view (Figure 2B, labels 1 and 2). In the “Result Display” page, positively predicted residues are highlighted in red in the sequence view (Figure 2C, label 1), and residue-level scores are shown in an interactive scatter plot (Figure 2C, label 2). Adjusting the threshold (Figure 2C, label 3) updates both the plot and the highlighted residues, but does not modify the downloaded prediction file. For LINKER-Pred2, users select the LINKER-Pred2 tab and submit a single FASTA sequence together with an .a3m MSA file. The remaining steps are identical to LINKER-Pred-Lite.

Downloads are provided as a ZIP archive named [TASK_ID]_results.zip, containing one CSV file per sequence ([sequence_name].csv). Each CSV file contains four columns: residue index (1-based), amino-acid letter, predicted linker score (0–1), and the binary label computed from the threshold selected at submission.

## CONCLUSIONS

Here, we present LINKER-Pred, a public web server for residue-level prediction and visualization of disordered flexible linkers (DFLs). The platform implements two convolutional ensemble predictors, LINKER-Pred2 (ProtTrans + MSA-Transformer) and LINKER-Pred-Lite (ProtTrans only), trained on curated DLD and DisProt linker datasets. Methodologically, the predictors validate the effectiveness of the compact M2M CNN architecture coupled to protein language model embeddings and ensemble averaging as an effective learning strategy for DFL prediction.

Linker prediction was one of the challenges showing the strongest improvement In CAID3, with respect to CAID 2, with ∼10% gain in AUC scores and ∼30% gain in APS among the top predictors [10,11]. This improvement was driven largely by new predictors, many of which were DFL-specific methods that implemented strategies such as pLM embedding, with LINKER-Pred scoring amongst the top-performing methods in CAID3. This underscores the growing impact of pLMs [20,29–31] in advancing intrinsic disorder prediction The refined LINKER-Pred predictors presented here were evaluated on the CAID2Linker (primary) and CAID3Linker (independent) benchmarks. The models achieve state-of-the-art AUC and APS across both benchmarks, with full coverage on all test proteins, demonstrating strong predictive power and practical robustness. On the independent CAID3 benchmark, the refined models outperform the original methods, and LINKER-Pred2 and LINKER-Pred achieve the highest AUCs among evaluated methods, with APS competitive with the leading predictors. Amongst other top-performing methods are IPA-AF2-Linker [24], which utilizes a CNN [32] trained on DisProt data, incorporating relative solvent accessibility (RSA) and prediction accuracy scores (pLDDT) from AlphaFold2 [22], and DisoFLAG-IDR and DisoFLAG-DFL [33], which employ a Graph-based Interaction Protein Language Model (GiPLM) trained on DisProt DFLs. However, the available benchmark datasets remain relatively small, and linker prediction is still an emerging field. Therefore, numerical differences between leading methods should be interpreted cautiously. Overall, the results support LINKER-Pred methods as competitive and practically robust predictors rather than establishing universal superiority over other approaches.

The modular architecture of the LINKER-Pred Web Server enables asynchronous, fault-tolerant, and scalable operation, supporting multiple concurrent users and large batch predictions without performance degradation. Computational components are containerized using Docker, which encapsulates dependencies and models to guarantee reproducibility. The web server deploys LINKER-Pred2 for accuracy-focused, single-sequence analyses and LINKER-Pred-Lite for rapid, multi-sequence throughput. This pairing delivers state-of-the-art accuracy where needed and efficient, scalable screening for routine and large-scale use.

Future work will expand training data to additional linker types and contexts, including candidate linker regions derived from large-scale AlphaFold-based structural analyses, and refine embedding strategies. We will also extend interoperability through batch APIs and workflow integration by incorporating the MSA generation into the pipeline. We will also explore optional sliding-window smoothing to convert residue-level scores into continuous linker-region predictions, facilitating interpretation of predicted linker boundaries and complementary segment-overlap evaluations against annotated regions. These advances will help further improve accuracy and throughput, consolidating the LINKER-Pred web server as a reliable resource for DFL prediction and for investigating protein modularity and functionality.

## AUTHOR ROLES

DM: Investigation, Formal analysis, Data curation, Writing – original draft; HMGA: Formal Analysis, Investigation, Methodology, Software, Validation, Visualization; JG: Formal analysis, Data curation, Investigation, Visualization, Writing – original draft, Supervision; COL: Resources, Software; GP: Supervision, Funding acquisition; LBC: Conceptualization, Supervision, Funding acquisition, Project administration, Writing – original draft. All authors reviewed and edited the final manuscript.

## DATA AVAILABILITY

The LINKER-Pred web server is publicly available at https://linkerpred.chemeslab.org/.

The training and validation datasets used in this study are available for download from the website’s **Download** section and are also provided in **Supplementary File S1**. The CAID Linker benchmark datasets used for model evaluation are provided in **Supplementary File S2**.

The full DLD dataset is available at https://dld.chemeslab.org/.

### GitHub

Embedding: https://github.com/deemeng/embedding

LINKER-Pred: https://github.com/deemeng/punch_linker

LINKER-Pred2: https://github.com/deemeng/punch_linker2

LINKER-Pred-Lite: https://github.com/deemeng/punch_linker_light

### Docker Images

Embedding: https://hub.docker.com/r/dimeng851/embedding

LINKER-Pred: https://hub.docker.com/r/dimeng851/punch_linker

LINKER-Pred2: https://hub.docker.com/r/dimeng851/punch_linker2

LINKER-Pred-Lite: https://hub.docker.com/r/dimeng851/punch_linker_light

