## SUPPLEMENTARY INFORMATION for "LINKER-Pred: A Deep Learning Method and Web-Server for the Prediction of Disordered Flexible Linkers in Proteins"

**This PDF file includes:**

Legends for Supplementary Files 1 and 2

Figure S1 to S3

Tables S1 to S6

Supplementary Note

Supplementary References

**Supplementary File S1 (Separate File). Training and validation datasets used for model development.** Datasets include protein sequences, residue-level target labels, source-dataset labels, cluster assignments, and cross-validation fold identifiers; each row represents one sequence, with target vectors aligned from the N- to C-terminus. Three partitioning schemes are provided: *Clustering_20%_ID* and *Clustering_30%_ID*, which use the indicated sequence-identity threshold both to remove CAID2-related sequences and to assign the remaining sequences to cluster-based cross-validation folds; and *Random_split*, which uses 20% sequence-identity filtering followed by random fold assignment.

**Supplementary File S2 (Separate File). CAID2Linker and CAID3Linker benchmark datasets.** Datasets include DisProt identifiers, protein sequences and residue-level target labels; each row represents one sequence, with target vectors aligned from the N- to C-terminus.

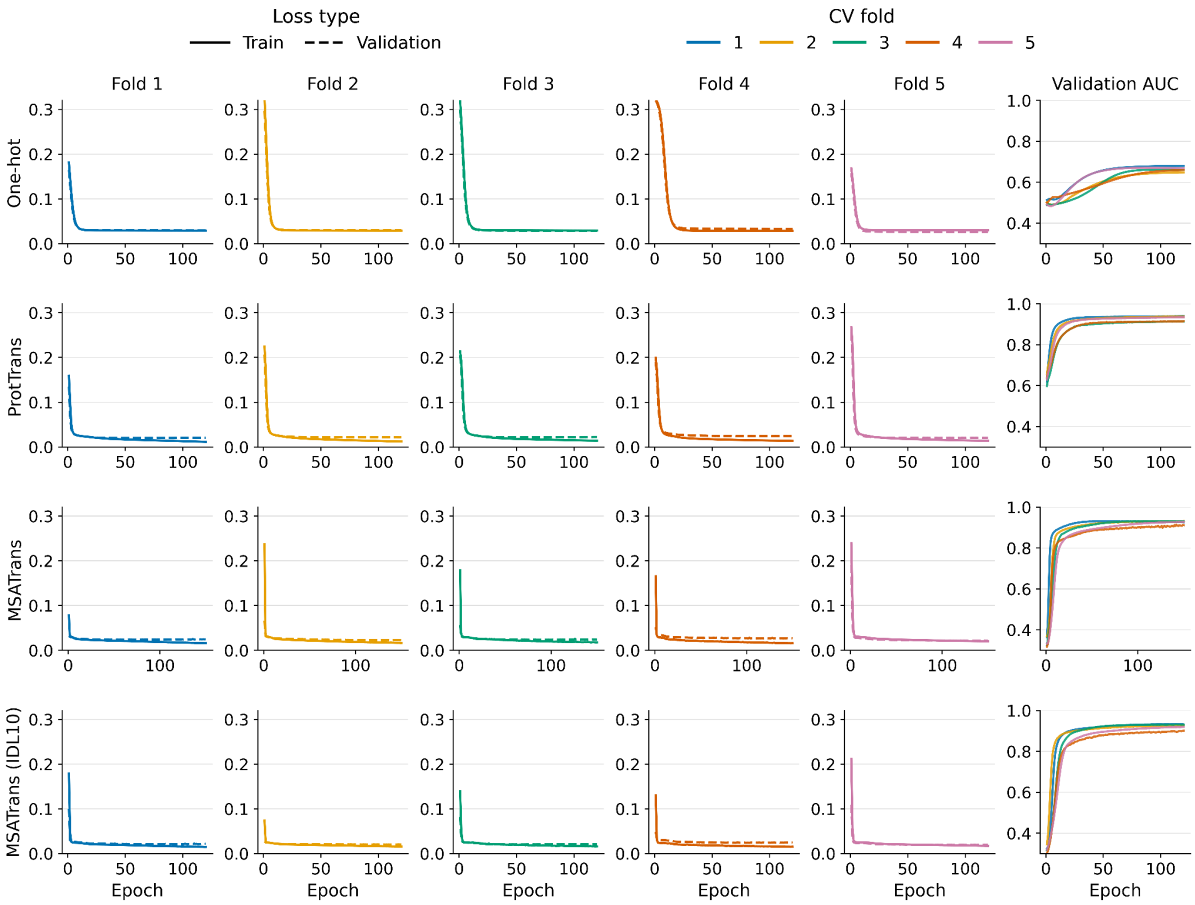


**Figure S1. Training histories across cross-validation folds using cluster-based partitioning at a 20% sequence-identity cutoff.** Training and validation loss are shown separately for each of the five folds with solid and dashed lines, respectively. The one-hot, ProtTrans, and MSA-Transformer input representations were trained using the DLD_IDL + DisProtDFL datasets, whereas the MSA-Transformer (IDL10) model was trained using DLD_IDL10 + DisProtDFL. The rightmost panels show validation ROC-AUC across training epochs, with colours distinguishing the five folds. Loss panels share the same y-axis range, while validation ROC-AUC panels share a separate common y-axis range, enabling comparisons across folds and model configurations.

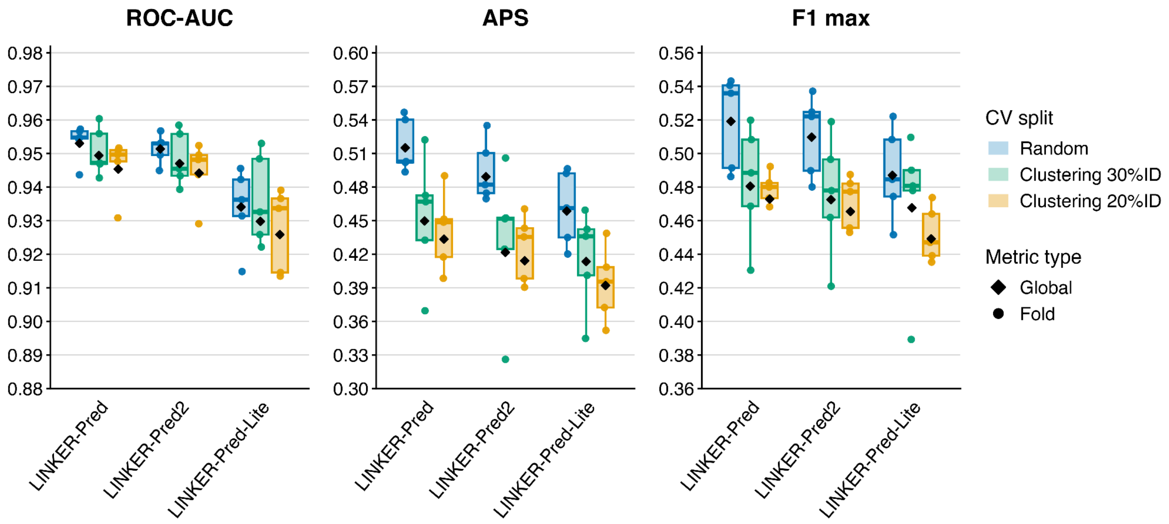


**Figure S2. 5-fold cross-validation performance of LINKER-Pred ensemble predictors.** Performance is shown for the three ensemble predictors trained with random and cluster-based data partitions. Boxplots summarize the distribution of fold-level metric values, with individual folds shown as circles. Black diamonds indicate the corresponding global metric, computed by pooling held-out residue-level predictions across the five folds. Colours indicate the cross-validation split strategy. Metrics are shown in separate panels: ROC-AUC, average precision score (APS), and maximum F1 score (F1 max). The y-axis range is scaled independently for each metric to improve visualization of within-metric differences and should not be compared across panels. Boxplots: centre line inside the box indicates the median value, box covers the interquartile range, and whiskers extend to, at most, 1.5-fold of the interquartile range.

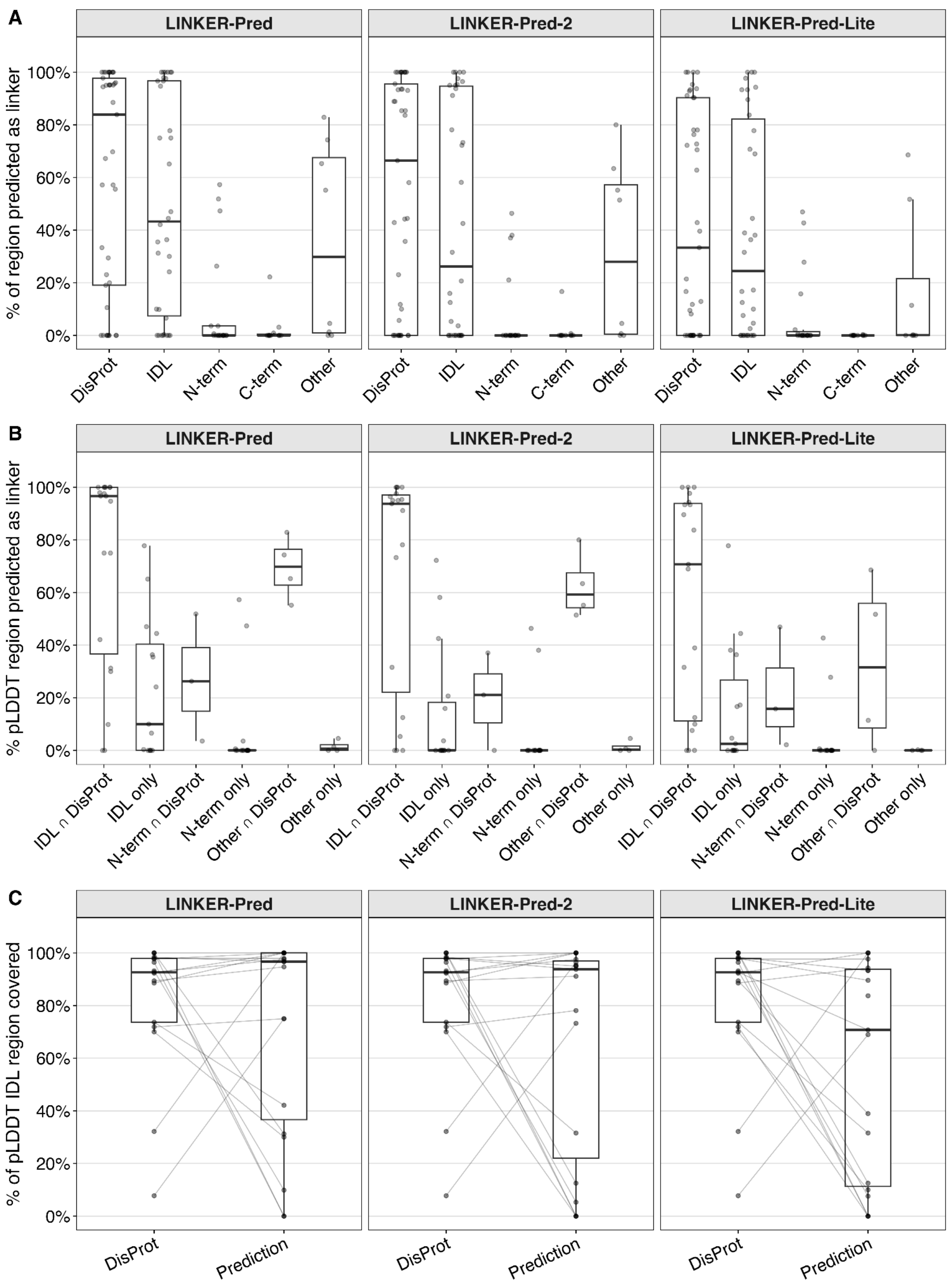


**Figure S3. Specificity of LINKER-Pred predictions across DisProt-annotated and pLDDT-derived low-confidence regions.** (A) Prediction coverage across DisProt DLFs (n=38) and pLDDT-derived regions classified as Inter-Domain Linkers (IDL, n=34), N-terminal regions (n=18), C-terminal regions (n=11), or Other internal regions (n=8). (B) Prediction coverage across pLDDT-derived regions stratified according to overlap with DisProt DLFs. Sample sizes: IDL ∩ DisProt, n = 19; IDL only, n = 15; N-term ∩ DisProt, n = 3; N-term only, n = 15; Other ∩ DisProt, n = 4; Other only, n = 4. C-term regions are absent from this panel since none overlapped a DisProt DFL. (C) For pLDDT-derived IDL regions overlapping DisProt DLFs (n=19), the % of the pLDDT IDL region covered by DisProt annotation is compared with the % predicted as linker by each LINKER-Pred method. Points represent individual regions. Boxplots: centre line inside the box indicates the median value, box covers the interquartile range, and whiskers extend to, at most, 1.5-fold of the interquartile range.

**Table S1: Overview of the training and benchmarking sets.**

| Category | Dataset | Source | Linker Type | # Total Seq | # Total DFLs | Description |
|---|---|---|---|---|---|---|
| Training | **DLD_IDL** | DLD | IDL | 2646 | 1640 | Entire DLD dataset |
| | **DLD_IDL10** | DLD | IDL | 2646 | 914 | Entire DLD dataset with linker annotations restricted to a minimum length of 10 |
| | **DisProtDFL** | DisProt | All types | 267 | 338 | DisProt sequences containing various linkers |
| **Primary Benchmarking** | **CAID2Linker** | DisProt | All types | 40 | 42 | Linker dataset from the CAID2 challenge |
| **Independent Benchmarking** | **CAID3Linker** | DisProt | All types | 31 | 37 | Linker dataset from the CAID3 challenge |

For DLD datasets, the number of total sequences (“#Total Seq”) refers to PDB chains, while for DisProt datasets it refers to protein sequences with annotated experimental evidence. “# Total DFLs” indicates the total number of annotated Disordered Flexible Linker regions. DLD: Domain-Linker-Domain; DFL: Disordered Flexible Linker; IDL: Independent Domain Linker.

**Table S2: Training sets after sequence identity filtering against CAID2Linker.**

| Category | Dataset | # Total Seq | #Remaining Seq | |
|---|---|---|---|---|
| | | | 30% ID | 20% ID |
| Training | DLD_IDL | 2646 | 2608 | 2573 |
| | DisProtDFL | 267 | 248 | 237 |
| **Combined Total** | | **2913** | **2856** | **2810** |

To prevent information leakage, sequences from the training sets showing sequence identity to any sequence in the primary benchmarking set (**CAID2Linker**) above the indicated cutoff were excluded. "# Total Seq" lists the original number of sequences in each dataset, whereas the "#Remaining Seq" columns indicate the number of sequences retained after filtering at 30% or 20% sequence identity. DLD_IDL10 uses the same DLD sequence set as DLD_IDL and is therefore not listed separately.

**Table S3: Overview of the training datasets, embeddings, and model configurations used in LINKER-Pred predictors.**

| Predictor | Dataset | | | Embedding | |
|---|---|---|---|---|---|
| | DLD_IDL | DLD_IDL10 | DisProtDFL | ProtTrans | MSATrans |
| LINKER-Pred (10 models) | X | – | X | 5 | – |
| | X | – | X | – | 5 |
| LINKER-Pred2 (15 models) | X | – | X | 5 | – |
| | X | – | X | – | 5 |
| | – | X | X | – | 5 |
| LINKER-Pred-Lite (5 models) | X | – | X | 5 | – |

An "X" indicates that a given dataset was used for training, while a "–" signifies that the dataset was not used. The numbers under the “Embedding” columns indicate the number of individual models trained for each dataset–embedding combination. Each value is 5 because individual models were trained using five-fold cross-validation. LINKER-Pred-Lite includes five ProtTrans-based models, LINKER-Pred includes five ProtTrans-based and five MSA-Transformer-based models, and LINKER-Pred2 additionally includes five MSA-Transformer-based models trained with the DLD_IDL10 and DisprotDFL datasets.

**Table S4. Composition of the final combined training sets used for individual model training.**

| Combined Datasets | CAID2 Seq ID Cutoff | # Seq Used | #Pos Seq | # Neg Seq | #DFLs |
|---|---|---|---|---|---|
| DLD_IDL + DisProtDFL | 30% | 2856 | 1630 | 1226 | 1897 |
| DLD_IDL10 + DisProtDFL | 30% | 2856 | 1063 | 1793 | 1177 |
| DLD_IDL + DisProtDFL | 20% | 2810 | 1596 | 1214 | 1858 |
| DLD_IDL10 + DisProtDFL | 20% | 2810 | 1036 | 1774 | 1147 |

Training sequences similar to the CAID2Linker benchmark were removed using the indicated sequence-identity cutoff. "# Seq Used" shows the number of sequences retained after sequence similarity removal. Positive sequences contain at least one DFL, whereas negative sequences contain no DFLs. "# DFLs" indicates the number of annotated disordered flexible linker regions in each combined training set.

**Table S5. Maximum F1 scores and corresponding optimal classification thresholds on the CAID2Linker and CAID3Linker datasets.**

| | CAID2Linker Dataset | | CAID3Linker Dataset | |
|---|---|---|---|---|
| **Predictor** | **F1 max** | **Best F1 threshold** | **F1 max** | **Best F1 threshold** |
| **LINKER-Pred2** | 0.316 | 0.080 | 0.546 | 0.130 |
| **LINKER-Pred** | 0.309 | 0.060 | 0.523 | 0.110 |
| **LINKER-Pred-Lite** | 0.289 | 0.060 | 0.434 | 0.150 |

Maximum F1 scores and corresponding optimal classification thresholds for the LINKER-Pred predictors on the CAID2Linker and CAID3Linker datasets. For each predictor and benchmark dataset, F1 max denotes the highest F1 score obtained across all evaluated decision thresholds, and the best F1 threshold indicates the threshold at which this maximum was achieved.

**Table S6. Results from CAID competition round 3**

| Predictor | AUC | APS | F1 max | Cov |
|---|---|---|---|---|
| IPA-AF2-Linker | 0.897 | 0.474 | 0.483 | 0.903 |
| **LINKER-Pred2** | **0.875** | **0.365** | **0.46** | **1** |
| **LINKER-Pred** | **0.87** | **0.377** | **0.471** | **1** |
| DisoFLAG-IDR | 0.866 | 0.312 | 0.444 | 0.967 |
| **LINKER-Pred-Lite** | **0.854** | **0.393** | **0.446** | **1** |
| DisorderUnetLM | 0.851 | 0.257 | 0.376 | 1 |
| DisoFLAG-DFL | 0.846 | 0.392 | 0.44 | 0.967 |
| APOD | 0.843 | 0.321 | 0.395 | 1 |
| flDPnn3a | 0.841 | 0.228 | 0.383 | 1 |
| ESMDisPred-2PDB | 0.836 | 0.345 | 0.443 | 0.806 |

The performance of our three linker predictors (LINKER-Pred, LINKER-Pred2, and LINKER-Pred-Lite) reported in CAID3 [1] differs slightly from that presented in Table 2. This discrepancy arises because, for the current study, we retrained all three models using an updated, higher-quality version of the DLD dataset. The improvements to the DLD dataset include: (a) exclusion of low-resolution PDB structures (resolution > 5 Å); (b) removal of short structural domains (<30 residues) following a two-step smoothing of short, structured elements; and (c) an increased inter-domain distance threshold (Cα–Cα) from 4.663 Å to 5 Å for distinguishing dependent and independent domains. Apart from this, instead of splitting the training data randomly, we employed a sequence-identity-based partitioning scheme (for more details, refer to Materials and Methods). Reported metrics are ROC-AUC, average precision score (APS), and maximum F1 score (F1 max). Finally, "Cov" refers to the sequence coverage or fraction of the CAID benchmark datasets that was predicted with each of the listed methods.

### Supplementary Note

#### Web server back end

The backend of the LINKER-Pred web server forms the computational and logical core of the platform. It handles web routing, user input, task creation, job execution, and file management through a Django-based framework [2]. The backend executes all major processes, including task validation, Docker control, and data tracking. When a sequence is submitted, Django's request handler validates the input format (FASTA and optional .a3m file), assigns a unique task ID, and generates a dedicated working directory containing the input, embedding, and output folders. Each task is then submitted to a Celery queue [3], which manages background execution without blocking the web interface.

Execution is controlled by a custom shell script that launches Docker containers for embedding and model inference. Separate images are used for sequence embedding, inference, and result packaging. This container-based design simplifies deployment and allows for consistent execution across environments. Depending on the input type, the system automatically selects either the LINKER-Pred-Lite or LINKER-Pred2 container image. After execution, containers are automatically removed, and the prediction outputs are stored in the designated result folder. For every submission, the backend logs detailed metadata including Task ID, creation time, prediction threshold, input/result file paths, and Task status. This database-driven structure allows users to retrieve completed results using their Task IDs. The system also includes cleanup utilities that automatically remove obsolete files, preventing storage overload and maintaining system stability.

#### Implementation and server runtime

The web server is implemented in Python 3.12 using Django for the web framework, Celery for task scheduling, and Redis as the message broker. When a user submits a prediction, Django dispatches a Celery task that triggers the corresponding Docker workflow. The container performs embedding and prediction, writes results to mounted directories, and exits automatically upon completion. All official Docker images are publicly available via Docker Hub. The system runs on a high-performance workstation equipped with an AMD Ryzen Threadripper 7960X (24 cores/48 threads) CPU, an NVIDIA RTX 4000 Ada GPU, and 256 GB RAM, operating under a 64-bit Linux system. Both LINKER-Pred and LINKER-Pred-Lite are GPU-accelerated but can also operate efficiently on CPUs alone.

We benchmarked the runtime performance of LINKER-Pred-Lite using 20420 human protein sequences from UniProtKB [4] (downloaded on 18 October 2025; 11,413,587 amino acids). The full dataset (.fasta) was submitted through the LINKER-Pred web server. The job finished in 20356 seconds, yielding an average processing speed of 1.78 seconds per 1000 residues, including embedding and inference, demonstrating that LINKER-Pred-Lite can handle proteome-scale analyses directly through the web interface without the need for local computation. LINKER-Pred2 performance was benchmarked against the CAID3Linker dataset using precomputed MSA files in .a3m format by submitting each sequence individually through the LINKER-Pred web server. The average runtime per prediction was 21.26 seconds, with an average of 32.15 seconds per 1000 residues, including embedding and inference. We did not benchmark MSA file generation, which was performed externally. Model inference itself remained comparable in speed to LINKER-Pred-Lite.

Overall, the LINKER-Pred web server combines an intuitive front-end interface with a scalable, containerized back-end. Users can submit sequences, monitor task progress, and interactively visualize prediction results through responsive web pages. All computations run asynchronously on a high-performance server, ensuring responsiveness under heavy workloads. Together, these components make the LINKER-Pred web server a robust platform for both exploratory and large-scale applications in protein structural bioinformatics.
